# Ancestral genome estimation reveals the history of ecological diversification in *Agrobacterium*

**DOI:** 10.1101/034843

**Authors:** Florent Lassalle, Rémi Planel, Simon Penel, David Chapulliot, Valérie Barbe, Audrey Dubost, Alexandra Calteau, David Vallenet, Damien Mornico, Thomas Bigot, Laurent Guéguen, Ludovic Vial, Daniel Muller, Vincent Daubin, Xavier Nesme

## Abstract

Horizontal gene transfer (HGT) is considered as a major source of innovation in bacteria, and as such is expected to drive adaptation to new ecological niches. However, among the many genes acquired through HGT along the diversification history of genomes, only a fraction may have actively contributed to sustained ecological adaptation. We used a phylogenetic approach accounting for the transfer of genes (or groups of genes) to estimate the history of genomes in *Agrobacterium* biovar 1, a diverse group of soil and plant-dwelling bacterial species. We identified clade-specific blocks of co-transferred genes encoding coherent biochemical pathways that may have contributed to the evolutionary success of key *Agrobacterium* clades. This pattern of gene co-evolution rejects a neutral model of transfer, in which neighbouring genes would be transferred independently of their function and rather suggests purifying selection on collectively coded acquired pathways. The acquisition of these synapomorphic blocks of co-functioning genes probably drove the ecological diversification of *Agrobacterium* and defined features of ancestral ecological niches, which consistently hint at a strong selective role of host plant rhizospheres.

## Introduction

Our understanding of bacterial ecology is fragmentary. We usually know a subset of the environments from which species can be sampled, a few laboratory conditions in which they can be grown, and sometimes the type of interactions they establish with other organisms. Their genomes, believed to encode all the information that make their lifestyle possible, are now available. However, even if we succeeded in describing the molecular function of each single base in a genome, we would not necessarily know whether this function is significant in the prevalent environment of the organism (Doolittle 2013). In order to discover those functions that are ecologically relevant, an approach consists in recognizing traces of selection for functional adaptation in the histories of genomes. Comparing genomes reveals historical signals that can be used to retrace genome evolution, by estimating their hypothetical ancestral state and the course of the evolutionary events that shaped them over time. Using models of null expectation under neutral evolution, we can discern the decisive events in the adaptive evolution of species.

Bacterial genomes are in constant flux, with genes gained and lost at rates that can exceed the nucleotide substitution rate (Lawrence & Ochman 1997). Recognition of this dynamics led to the concept of pangenome, i.e. the set of all homologous gene families present in a group of related genomes. The pangenome is the sum of the core and accessory genomes, which respectively gather the genes shared by all genomes in the dataset and those found in some genomes only. In *E. coli,* for example, the core genome is estimated to include 1,800 gene families, while the accessory genome has more than 80,000 gene families, with two random strains typically differing by a thousand (Touchon et al. 2009; Land et al. 2015). In a genome, accessory genes are regularly gained (notably by transfer) or lost, leaving patterns of presence in genomes that are inconsistent with the strain phylogeny (Young 2016).

For a majority of accessory gene families, this process occurs so rapidly that they are effectively observed in a single genome, caught by the snapshot of genome sequencing. This suggests that they only have transient, if any, adaptive value for their bacterial host (Daubin et al. 2003). However, this constant input of genes also allows adaptive accessory genes to settle in genomes, and become part of the core genome of a lineage. Such ‘domestication’ events amidst the rapid turnover of genome gene content constitute the most remarkable deviations from a neutral model in which all genes are equally likely gained and lost. Clade-specific conservation of a gene is thus suggestive of adaptation to a particular ecological niche (Lassalle et al. 2015).

In a previous study, we investigated the diversity of gene repertoires among strains of *Agrobacterium* biovar 1 (Lassalle et al. 2011). This taxon contains several bona fide yet unnamed ‘genomic’ species, numbered G1 to G9 and G13 and collectively named ‘*Agrobacterium tumefaciens* species complex’ (*At*) according to the proposal of Costechareyre et al. (Costechareyre et al. 2010). Genes specific to the species under focus *–* G8, for which we proposed the name *A. fabrum* – were usually physically clustered in the genome, and these clusters in turn gathered genes that encoded coherent biological functions (Lassalle et al. 2011). The conservation of co-functioning genes in genomic clusters appears unlikely in the context of frequent gene turnover. This pattern could be a trace of purifying selection that led to retain the whole gene clusters, because the selected unit was the function collectively encoded by the constituent genes. However, it could also result from a neutral process of gene flow, whereby neighbour genes with related functions (e.g. operons) happen to be transferred together and are then slowly eroded. These hypotheses may however be distinguished by analysing the historical record of evolutionary events that led to the clustering of co-functioning genes.

Most genes have complex histories, marked by many events of gene duplication, loss and, especially in the case of micro-organisms, horizontal transfers. The set of events affecting each homologous gene family in the pangenome under scrutiny can be summarized into an evolutionary scenario that can be seen as the path of gene evolution within and across branches of the tree of species. Evolutionary scenarios can be inferred by comparing the phylogenetic history of genes to the phylogenetic history of species, and by reconciling their discordances through the explicit inference of duplication, transfer and loss events (Doyon et al. 2011; Scornavacca et al. 2012). This in turn makes it possible to deduce the incremental shaping of genome gene contents, from ancestral to contemporary genomes, and to try and deduce the functional and ecological consequences of these changes.

We used the Rhizobiaceae family as a model taxon, and more particularly focused on the *At* clade for which we gathered a dataset of 22 strain genomes from ten different species, including 16 newly sequenced genomes. We designed a new phylogenetic pipeline for the estimation of ancestral genome gene contents that accounts for horizontal gene transfer and gene duplication. Applied to our dataset, this approach estimated blocks of co-transferred and co-duplicated genes, enabling us to test hypotheses on how co-functioning gene clusters were formed. Then we compared the level of functional co-operation of genes within blocks of co-transferred clade-specific genes to the expectation under a neutral model of horizontal gene transfer where genes are randomly picked from the donor genome. This comparison showed that clade-specific genes were more functionally related than expected, supporting the hypothesis that domestication of at least some clade-specific genes resulted from ecological selection.

Our estimated pangenome history – from single gene trees with transfer and duplication events to blocks of co-evolved genes and functional annotations – was compiled in an integrative database called Agrogenom, which can be visualized and queried through an interactive web interface accessible at http://phylariane.univlyon1.fr/db/agrogenom/3.

## Methods

### Bacterial culture experiments

Bacterial growth was analysed in the presence of phenylacetate (5mM) using a Microbiology Bioscreen C Reader (Labsystems, Finland) according to the manufacturer’s instructions. Pre-cultures of *Agrobacterium* strains were grown overnight in AT medium supplemented with succinate and ammonium sulphate. They were inoculated at an optical density at 600 nm (OD_600_) of 0.05 in 200 µl AT medium supplemented with appropriate carbon and nitrogen sources in Bioscreen honeycomb 100-well sterile plates. Cultures were incubated in the dark at 28°C for 3 days with moderate shaking. Growth measurements (OD_600_) were performed at 20-min intervals.

#### Genome sequencing and assembly

Genomic DNAs of the 16 *At* strains (Table 1) extracted with the phenol-chloroform method were used to prepare libraries with DNA sheared into 8-kb inserts (median size). Raw sequence data were then generated using 454 GS-FLX sequencer (Roche Applied Sciences, Basel, Switzerland) with a combination of single-read (SR) and mate-pair (MP) protocols that yielded coverage ranging from 6.5X to 11X and from 5X to 8X, respectively (Table S1). Genome sequences were then assembled with Newbler version 2.6 (Roche Applied Sciences, Basel, Switzerland), using 90% identity and 40-bp thresholds for alignment of reads into contigs and the ‘--scaffold’ option to integrate duplicated contigs into the scaffold assembly. Virtual molecules (chromosomes and plasmids) gathering scaffolds were manually created on the basis of plasmid profiles obtained from Eckhart gels (data not shown) and minimizing rearrangements between closely related genomes by taking into account whole-genome alignments obtained with the NUCmer program from the MUMMER package version 3.0 (Kurtz et al. 2004). Genome sequences were then annotated with the MicroScope platform (Vallenet et al. 2013) and made available through the MaGe web interface (www.genoscope.cns.fr/agc/microscope) or the European Nucleotide Archive (http://www.ebi.ac.uk/ena/data/view/<ACCESSION NUMBERS> with accessions starred in Table 1).

**Table 1:**
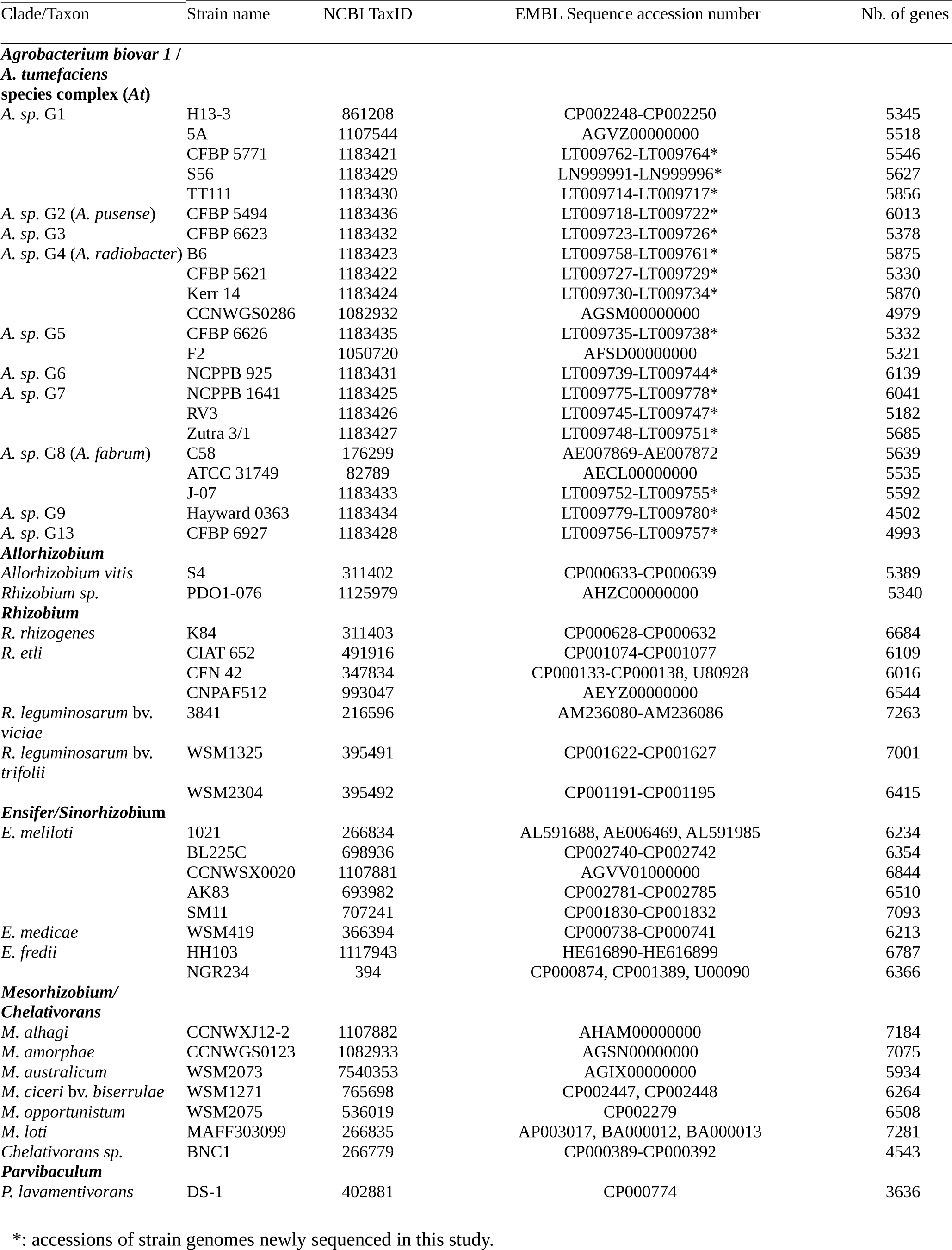
List of the 47 Rhizobiales strains used in this study.

#### Genomic sequence dataset

The study focused on the *Agrobacterium* biovar 1 species complex a.k.a. *A. tumefaciens* (*At*) with an original dataset of the aforementioned 16 new genomes, plus six publicly released ones (Goodner et al. 2001; Wood et al. 2001; Li et al. 2011; Ruffing et al. 2011; Wibberg et al. 2011; Hao, Lin, et al. 2012; Hao, Xie, et al. 2012). These 22 genomes covered 10 closely related but genomically differentiated species (G1 to G9 and G13), with up to five isolates per species. The dataset also included all Rhizobiaceae genome publicly available at the time of the database construction (spring 2012), and a few more distant relatives from the Phyllobacteriaceae and Rhodobiaceae families (Table 1; Fig. S1).

#### Homologous gene family database

Based the 47 complete genome sequence dataset, we built a database of homologous gene families following the model of Hogenom databases (Penel et al. 2009). All annotated protein coding sequences (CDSs) were extracted and translated into protein sequences on which a all-versus-all pairwise BLASTP similarity search was performed to build a similarity network. Homologous gene families were derived from the connected components of the network using HiFix (Miele et al. 2012). Gene family sequences were then aligned at the protein level using MUSCLE (Edgar 2004) and reverse-translated into CDS alignments with pal2nal (Suyama et al. 2006). For an extended description of bioinformatic procedures, please refer to Supplementary Text, section 2.

#### Reference species tree

To construct the reference species tree, we used 455 unicopy core gene families (i.e. families with exactly one copy per genome, listed Table S2), proceeding to 500 jackknife samples (draws without replacement) of 25 gene alignment sets, which were each concatenated and used to infer a maximum-likelihood (ML) tree using PhyML (Guindon & Gascuel 2003) using the same parameters as for gene trees (see Supplementary Text). The reference phylogeny was obtained by making a consensus of this 500-tree sample with the CONSENSE algorithm from the Phylip package (Felsenstein 1993), and branch supports were derived from the frequency of the consensus tree bipartitions in the sample (Fig. S2). Alternative phylogenies were searched using the concatenate of the whole set of 455 universal unicopy families or from a concatenate of 49 ribosomal protein gene families (Table S3) to compute trees with RAxML (version 7.2.8, GTRCAT model, 50 discrete site-heterogeneity categories) (Stamatakis 2006).

#### Reconciliation of genome and gene tree histories

We computed gene trees using PhyML (Guindon & Gascuel 2003) for all 10,774 gene families containing at least three genes (see Supplementary Text, section 3) and estimated the branch support using the SH-like criterion. We rooted these gene trees using the combo criterion of TPMS (Bigot et al. 2013) so that, knowing the species phylogeny, both species multiplicity and taxonomic depth of all subtrees were minimized. A root minimizing these criteria favours reconciliation scenarios with less ancient gain (duplication and transfer) event, leading to scenarios more parsimonious in subsequent losses (Fig. S3, step 1). As this criterion yields poor results in the absence of ancestral duplications and the presence of many transfers, we used another method to root unicopy gene trees (i.e. trees of gene families with one gene per genome at most): we ran Prunier (Abby et al. 2010) for HGT detection (see below) and retained the root consistent with the most parsimonious transfer scenario.

We then inferred an evolutionary scenario for each gene family, i.e. a mapping in the species tree of the presence/absence of gene lineages and of the events that led to their emergence. We reconciled the gene tree topologies with the species tree by annotating each of the 467,528 nodes found in the 10,774 gene trees with an estimated event of origination, duplication, transfer (ODT), or speciation. We used a bioinformatic pipeline that combines several methods dedicated to the recognition of different signals of duplication and horizontal transfers, fully detailed in the Supplementary Text, Section 3, and summarized below and in Table S4. In brief, gene trees were processed as follows: likely duplication events were first located by looking for clades with multiple gene copies per species (Fig. S3, step 2). Within the implied paralogous clades, subtree pruning and regrafting (SPR) moves that did not disturb branches with high (≥ 0.9) support were attempted, and retained as topology updates when they decreased the incidence of duplication events (by reducing the count of events or the count of descendant gene tree leaves). Another 17,569 nodes remained marked as putative duplications, out of which 28,343 potential paralogous lineages emerged. We used those as guide to extract subtrees in which every species was represented once, i.e. unicopy subtrees. To deal with lineage-specific paralogues (‘in-paralogues’), we extracted the several possible combinations of co-orthologous gene copies (see Kristensen et al. 2011), producing unicopy subtrees with different but overlapping leaf sets (Fig. S3, step 3). Prunier, a parsimony-based method that takes into account the phylogenetic support of topological incongruences (Abby et al. 2010), was run on the unicopy subtrees to detect replacing transfer events based on significant topological conflict, i.e. involving branches with statistical support greater than 0.9 (Fig. S3, step 3). These reconciliations of potentially overlapping local subtrees yielded point estimate scenarios (involving a total of 22,322 phylogenetically supported transfer events), which were mapped back to the gene trees (Fig. S3, step 4). When several alternative (possibly conflicting) reconciliation scenarios were generated by independent inferences on overlapping lineage subtrees ("replicates"), the most likely scenario was chosen based on the number of similar events inferred in the neighbouring gene families (Fig. S3, step 5), favouring the events involved in the largest block events (see the “Block event inference” section below).

In the next step, we completed the reconciliation of gene tree topologies with the species tree topology: topological incongruences may still have remained, notably involving gene tree branches with statistical support too low for Prunier to identify them as significant topological conflicts and to propose a transfer event. These topological incongruences needed to be explained – notwithstanding branch supports – by scenarios involving duplications or transfers (and subsequent losses), transfer scenarios being usually more parsimonious in the count of invoked events. We thus used the taxonomic incongruence algorithm from Bigot et al. (2013) to identify 1,899 conflicting branches as the places of additional transfer events, where otherwise 10,229 additional counts of duplication events would have been necessary (Fig. S3, step 6). This gave us a final estimate of the collection of duplication and horizontal transfer events leading to the emergence of new gene lineages. We then defined a subfamilies of orthologues (nested in homologous gene families) as the descendants of every gene gain (ODT) event (Fig. S3, step 6). Finally, we used the Wagner parsimony algorithm implemented in the Count program (Csűrös 2008) to estimate scenarios of orthologous subfamily evolution, where transfers can be inferred to explain heterogeneous profiles of gene occurrence. This led to the annotation of 19,553 additional transfer events (Fig. S3, step 7). The illustrated description and programming details of the reconciliation pipeline used in this studies are available at: https://github.com/flass/agrogenom/blob/master/pipeline, and intermediary input/ouput files and datasets are available at: https://figshare.com/projects/Ancestral_genome_reconstruction_reveals_the_history_of_ecological_diversification_in_Agrobacterium/20894.

### Coordinates of origination, duplication and transfer events in the species tree

Transfer events are characterized by the location of both donor and receiver ancestor nodes in the species tree (further referred to as “event coordinates”), which specifies the direction of the transfer; other gene gain events – gene origination or duplication – are only characterized by their location at an ancestral node in the species tree. The inference of co-events (events that involved several genes, see “Block event inference” below) relies on the detection of similar events across gene families, i.e. events with the same coordinates. However, this can be challenging because independent evolution of gene families after a co-event may leave very different patterns in the respective gene trees, for instance due to different histories of gene loss after a common ancestral gain by co-transfer. When losses are considered, the right counts and locations of events are notoriously hard to estimate, as many combinations of loss events are possible for a fixed number of gain events, with little information – only gene absence, i.e. missing data – to inform a choice. To get a point estimate of a scenario with gains and losses, one typically applies a criterion of parsimony on the count of loss events subsequent to a gain (e.g. by transfer), so the gain event is estimated to be located at the last common ancestor of the species represented in the recipient clade. Given that gene families often have different species representations, this can result in family-specific systematic biases when estimating event coordinates. Unmatched biases in coordinate estimates would strongly affect our ability to recognize that a same co-event affected neighbour gene families (Fig. 1 A). To reach a higher sensitivity in detecting similar events, we left counts and locations of loss events undetermined. This resulted in degrees of freedom on the ODT scenarios, with several connected branches of the species tree on which ODT events could possibly have happened (Fig. 1 A). As a result, we represented ODT event coordinates as sets of species tree nodes; two such sets are necessary in the case of transfers to characterize both donor and recipient locations (Fig. 1 A, inset table).

**Figure 1:**
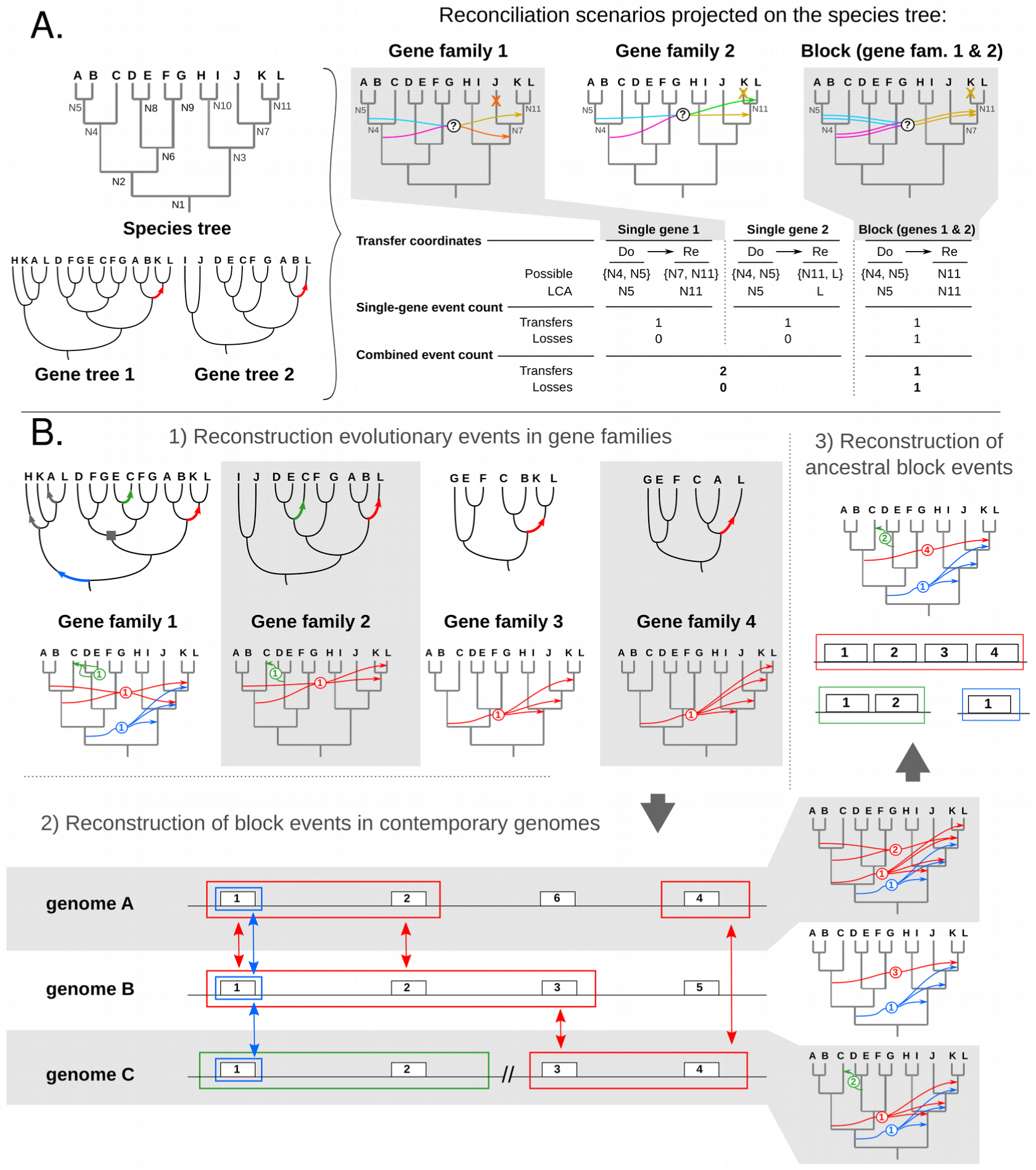
Single gene vs. block event reconciliation. A) Transfers inferred in reconciled gene trees 1 and 2 can be translated into several possible scenarios in the species tree, and each scenario involves different donor (Do) and receiver (Re) pairs (multiple arrows with question marks, uncertain scenarios). If each gene family is reconciled separately, the scenarios that place the ancestral receiver as the last common ancestor of extant recipient genomes were chosen because they were the most parsimonious in losses (crosses mapped on the species tree and “Local event count” in inset table). In that way, the global scenario for the combined loci totalizes two transfers and no subsequent loss (inset table, “Combined event count”). If the transfer event coordinates are compatible (i.e. non-null intersection: Re:{N7, N11} ∩ Re:{N11, L} = Re:{N11}) between gene families, we hypothesized the co-transfer of neighbour genes 1 and 2 as a common (Block) transfer event. By accounting for co-transfer events, a scenario was chosen which was not necessarily the most parsimonious one as regards losses for each gene. In this example, the most parsimonious global scenario for the combined loci totalled one block transfer and one subsequent gene loss. B) Scheme of block event estimation. Origination, duplication and transfer events were first estimated separately in each gene family (1); for the sake of clarity, the example shows only transfer events, represented as arrows on gene tree branches (top) and between species tree branches (bottom). Compatible events affecting genes that were neighbour in at least one extant genome were aggregated into blocks (coloured frames) (2) and this approach was then repeated across genomes (vertical double arrows) to estimate in which ancestral genomes the events occurred (3). Circled numbers indicate the number of genes combined into a same event.

#### Block event inference

We define block events as unique ODT events that involved a block of several contiguous genes in an ancestral genome (‘ancestral block event’); by extension, ‘leaf block events’ refer to the blocks of genes descended from such an ancestral block event, which typically form syntenic blocks in extant genomes and share a similar evolutionary pattern. We used a greedy accretion procedure that 1) linked matching events from neighbour gene families together into leaf block events, and 2) linked all homologous leaf block events to a common ancestral block event (Fig. 1 B). The complete algorithm for block event inference is described in the Supplementary Text, section 4, and summarized below.

##### Leaf block event inference

Using a greedy algorithm similar to that defined by Williams et al. (2012), we built leaf block events by iterative inclusion of events from contiguous gene families with compatible coordinates. For each replicon (chromosome or plasmid) in the database, we iterated over each gene following their position on the replicon; the nodes on the reconciled gene tree lineage leading to this gene were evaluated from tip to root. If a node was associated to an ODT event, we initiated a leaf block event containing this event as seed, and set the block coordinates as those of the seed ODT event. Then we looked for a similar event in the gene tree of the direct neighbour gene, using the same procedure to scan its lineage from tip to root. If the event associated to a node was of the same nature (O, D or T) and with compatible coordinates (Fig. 1 A), it was appended to the leaf block event; the coordinate set of the leaf block event was then refined as the intersection of the coordinate sets of the block event and of the newly added event. When a matching event was found, this iterative search was repeated on the next neighbour gene’s lineage. In spite of finding such matching event, a leaf block event was extendable with a maximum of *g* ‘gap’ genes (*g*_O_ = 1; *g*_D_ = 0; *g*_T_ = 4), and its elongation was terminated if no gene with a matching event was found beyond (Fig. S4 A, B).

In the particular case of transfer (T) events, after the termination of a leaf block, inner gap genes were checked for phylogenetic compatibility of their gene tree with the scenario associated to the leaf block event (Fig. S4 C): we checked that clades of donor and receptor species were not separated from each other in the gene tree by any strongly supported branches. When no branches or only branches with weak statistical support (< 0.9) separated the clade pair, the transfer event hypothesis was not rejected and the leaf block event integrity was maintained. Conversely, when the gene tree of a gap gene carried a strong signal rejecting the transfer event, the original leaf block was split into two leaf blocks representing separate transfer events (Fig. S4 D).

##### Ancestral block event inference

Then, we estimated ancestral block events by searching homology relationships between leaf block events. Block homology was defined as the presence in each leaf blocks of at least one homologous gene associated to the same gene tree event (Fig. 1 B, step 2); this relationship can be found between leaf block events from different extant genomes or from a same genome. Ancestral block events were iteratively assembled from homologous leaf block events, and their coordinates were estimated by intersecting the coordinates of their members (Fig. 1 B, step 3).

This last step notably united certain leaf block events scattered in an individual genome. This allowed us to infer the unity of ancient gene blocks that were larger than their derived forms in extant genomes. Because of gene insertion/deletion or genomic rearrangement, contiguity of genes descending from a same co-event could easily have been disrupted. Due to this mutational process, the gene content of putative homologous leaf block events could differ, and their estimated block event coordinates could differ too. The leaf block homology relationship is supposed to be transitive, but due to these potential differences, incompatibilities could arise during the iterative accretion of leaf block events into ancestral events; in that case a heuristic was used to resolve the conflict between putative homologous leaf block events and distribute them into a number of self-compatible ancestral blocks.

##### Detection of block events in Agrogenom scenarios

Block events were investigated for origination (O), duplication (D) and transfer (T) events. We did not investigate losses (L), because random convergent losses occur at a higher rate (Kuo et al. 2009; David & Alm 2011; Szöllősi et al. 2012), and the larger solution space of loss scenarios leads to a higher risk of non-specific aggregation of unrelated loss events. For a similar reason of a high risk of false positives, we did not investigate O and D block events on the deep, long branches of the species tree (Fig. S1: N1, N2 and N3, respectively leading to *Parvibaculum lavamentivorans*, the *Mesorhizobium*/*Chelativorans* clade and the Rhizobiaceae clade), where many events were annotated with indistinguishable coordinates that likely occurred separately over time (2,586 O events and 2,934 D events overlooked). After all homology search, the coordinates of the ancestral block events for O, D and T were finally reduced to their most recent possible location in the species tree and subsequent losses were inferred accordingly to complete the gene evolution scenarios (point estimates for each gene family).

#### Detection of clade-specific genes from phylogenetic profiles

Clade-specific genes were defined as genes gained (or lost) by the clade ancestor and conserved (not re-gained) in all clade members since. We first identified genes marked by gain/loss events in the genome of a clade ancestor. Then, we identified clade-specific genes by searching for contrasting patterns in the phylogenetic profile of the presence or absence of each gained/lost gene. These profiles were established from the scenarios of orthologous subfamily evolution (see above and Supplementary Text, section 3, step 6). A background clade was chosen as the one corresponding to the next higher taxonomic unit (genus, species complex, etc.) in which the focal (foreground) clade was nested. Contrast was initially defined between the foreground and background clades, where foreground genomes had a consistently opposite pattern to that of genomes in the background clade. However, possible subsequent transfer or loss events in the background clade can blur the contrasting pattern in phylogenetic profiles. Clade-specific genotypes were thus identified using a relaxed definition of clade specificity, i.e. where the presence/absence contrast could be incomplete, with up to two genomes in the background clade sharing the foreground state.

#### Functional homogeneity of gene groups

To measure to which extent co-transferred genes showed coherence in the functions they encoded, we used metrics of semantic similarities of the Gene Ontology (GO) terms annotated to the gene products. First, we retrieved GO annotations from UniProt-GOA (http://www.ebi.ac.uk/GOA/downloads, last accessed February 2nd, 2013) (Dimmer et al. 2011) for public genomes, and used a similar pipeline of association of GO terms to gene products to annotate the genomic sequences produced for this study. The results of several automatic annotation methods were retrieved from the PkGDB database (Vallenet et al. 2013) based on similarity searches: HMM profile searches on InterPro, HAMAP and PRIAM databases and BLASTP searches on the SwissProt and TrEMBL databases (as of the February 5th, 2013), with a general cut-off e-value of 10e-10. GO annotations were then mapped to gene products using mappings between those method results and GO terms as provided by Uniprot-GOA for electronic annotation methods (http://www.ebi.ac.uk/GOA/ElectronicAnnotationMethods, last accessed February 12th, 2013): InterPro2GO, HAMAP2GO, EC2GO, UniprotKeyword2GO, UniprotSubcellular_Location2GO. The annotation dataset was limited to the electronically inferred data to avoid biases in the annotation of certain model strains or genes. The resulting functional annotations of proteomes were analysed in the context of Gene Ontology term reference (full ontology file downloaded at http://www.geneontology.org/GO.downloads.ontology.shtml, last accessed September 2nd, 2013) (Ashburner et al. 2000). Functional homogeneity (*FH*) within a group of genes is defined as the average value of the pairwise functional similarities between all gene products in the group, each of which is the average value of pairwise similarities between all terms annotated to a pair of genes. Similarities were measured using the *Rel* (within a gene) metric and the *funSim* metric (between genes) (Schlicker et al. 2006; Pesquita et al. 2009). Computations were done using a custom Python package derived from AIGO package v0.1.0 (https://pypi.python.org/pypi/AIGO).

To assess if co-transfer of genes was associated with coherent functions, we compared the *FH* of co-transferred gene blocks to that of random groups of genes, obtained either by uniformly sampling (i.e. by random drawing without replacement) individual non-linked genes or by sampling genomic windows of neighbour (linked) genes. *FH* values were computed for all windows of neighbour genes around a replicon, and a sample of the same size was drawn for random combinations of non-linked genes. Because the size of the group of genes strongly impacts the computation of the similarity metrics, and because the annotation density can vary among organisms and replicons (contiguous DNA molecules), the distributions of *FH* values were calculated per replicon and per group size. Note that the set of blocks of co-transferred genes is included in the set of all genomic windows, but that we used non-overlapping subsets for statistical comparisons.

To test if functional coherence of a block of co-transferred genes impacted its probability of retention after transfer, we compared the *FH* values of genes from two sets of ancestral block events: those where all constituent genes were conserved in all descendant leaf block events, and those where part of the genes were lost in at least one descendant leaf block events. To avoid biases linked to variation in age of transfer events, this comparison was made only for events that occurred in ancestors of species-level clades of *At.*

#### Agrogenom database

All data about genes (functional annotations, gene families), genomes (position of genes, architecture in replicons …), the species tree (nodes, taxonomic information), reconciliations (gene trees, ODT events), block events, inference analyses (parameters, scores …), and all other data relative to the present work were compiled in a PosgreSQL relational database called Agrogenom. The database schema, input data and build procedure are available at https://github.com/flass/agrogenom/tree/master/pipeline/database; its content is browsable through a web interface at http://phylariane.univ-lyon1.fr/db/agrogenom/3/.

## Results and Discussion

### Genomic Dataset and Reference Species Tree

To explore the genomic diversity of the Rhizobiaceae pangenome, we gathered 47 genomes from the *Agrobacterium*, *Rhizobium*, *Sinorhizobium*/*Ensifer*, *Mesorhizobium*/*Chelativorans* and *Parvibaculum* genera into the Agrogenom database. These genomes contain 281,223 coding sequences (CDSs, or genes hereafter) clustered into 42,239 homologous gene families. Out of these families, 27,547 were singletons with no detectable homologues (ORFan families) and 455 were found in exactly one copy in all 47 genomes (unicopy core gene families). Following the procedure used in Abby et al. (2012), a species phylogeny was inferred from the concatenation of unicopy core gene family alignments, using jackknife re-sampling of genes to compute branch supports (Fig. S2). Significant support was obtained for all clades corresponding to previously described species: *S. melitoti, R. etli, R. leguminosarum*, and in particular *At* species G1, G8, G4, G5 and G7. In contrast, branch support was low for the relative positioning of most strains within species, showing conflicting (or a lack of) signal among concatenated genes. Within the *At* clade, higher-order groupings were also highly supported: G8 with G6 (hereafter named [G6-G8] clade), G5 with G13, ([G5-G13] clade), G1 with [G5-G13] ([G1-G5-G13] clade), G3 with [G1-G5-G13] ([G3-G1-G5-G13] clade), G7 with G9 ([G7-G9] clade), and G4 with [G7-G9] ([G4-G7-G9] clade). Only a few deep splits such as the position of species G2 and [G6-G8] clade relatively to the *At* root were poorly supported (Fig. S2). We compared this species tree topology to two others obtained with alternative datasets (see Methods): all three methods yielded very similar results concerning the placement of the different genera and species (Fig. S5); the main difference resided in the rooting of *At* within the Rhizobiaceae clade, and the placement of lone representatives for species G2 and G3. Investigation of the pangenome-wide support for alternative hypotheses (see Supplementary Text, section 1; Fig. S6) confirmed that the best topology was provided by the jackknife sample consensus tree presented Fig. S2. A phylogeny estimated from the genome gene contents proved less appropriate to discriminate species, indicating the occurrence of a large quantity of HGTs (Fig. S7).

### Reconciliation of gene and species histories

To estimate the history of HGT and other macro-evolutionary events that shaped the Rhizobiaceae pangenome, we reconciled the topologies of gene trees with the species tree, i.e. we explained their incongruence by assigning events of origination, duplication, transfer (ODT), or speciation to the gene tree nodes. We used a succession of heuristics for the reconciliation of gene and species trees aimed at solutions parsimonious in losses and transfers (Fig. S3). The combination of events estimated in each gene tree resulted in an estimated scenario of evolution of the gene family along the species tree.

Out of the 467,528 nodes found in the rooted gene trees of the 10,774 families that contained at least three genes, our pipeline assigned a total of 7,340 duplication events (1.5% of all gene tree nodes) and 43,233 transfers (9.2%). The remainder of unannotated gene tree nodes corresponded to speciation events (where gene tree topologies locally follow the species tree) and originations (emergence of the gene family in our dataset, mapped at the root of the gene tree) (Table 2). Based on the estimated ancestral genome gene contents, we distinguished additive transfers that brought new genes, as opposed to those that replaced current orthologous genes. Replacing transfers accounted for a quarter of total transfers (9,271 events). Additive transfers contribute almost five times more than duplications to the total gene input in genomes (Table 2), showing that transfer is the main source of gene content innovation in *At*.

**Table 2:** Origination, duplication, transfer and speciation events estimated in reconciliations of the Agrogenom database.

| Event type | Single gene events | Block events | (of size >1) | Difference after event integration into blocks |
| --- | --- | --- | --- | --- |
| Originations | 5,189 | 4,267 | (667) | -922 |
| Duplications | 7,340 | 5,819 | (778) | -1,521 |
| Total transfers | 43,233 | 32,255 | (5,649) | -10,978 |
| Replacing transfers † | 9,271 | - | - |  |
| Additive transfers † | 33,962 | - | - |  |
| Total ODT | 55,762 | 42,341 | (7,094) | -13,421 |
| Implied losses | 29,843 | 32,739 | - | +2,896 |
| Total ODTL | 85,605 | 75,080 | - | -10,525 |

O, origination; D, duplication; T, transfer; L, loss; ODT refers to the combination of all O, D and T events, while ODTL also includes losses. †: Replacing and additive transfers were not distinguished in block events.

### Identification of co-events involving neighbour genes leads to a more parsimonious genome-wide scenario

Large-scale comparative genomics analyses revealed that insertions in genomes are typically composed of several consecutive genes, indicating that blocks of genes can evolve in linkage across genomes (Vallenet et al. 2009). Yet, to date, gene evolution scenarios have generally been evaluated for each gene tree independently of its neighbours (Makarova et al. 2006; Kettler et al. 2007). This is questionable because a scenario may be optimal (e.g., more parsimonious) for a given gene, but sub-optimal in a model where genes can be part of the same event (Fig. 1). We developed a procedure to identify blocks of genes that likely co-evolved through the same event, based on the compatibility of their coordinates in the species tree (see Methods).

By assembling compatible ODT events from individual reconciliations of neighbour genes, we inferred putative ‘block events’, i.e. unique evolutionary events that involved blocks of neighbour genes (Fig. 1 B,C). At the pangenome scale, we found numerous such block events in *At* genomes, with 17.5% of transfers and 13.3% of duplications involving at least two genes (Table 2). Several thousands of transfer events were inferred to involve 2 to 6 genes, and a few hundreds to span a dozen or more consecutive genes in extant genomes (Fig. S8 A). Moreover, blocks of ancestral genes that we estimated to have been transferred among ancestral genomes (‘ancestral block events’) often appeared as larger units than their extant counterparts (Fig. S8 B), indicating that rearrangements and partial losses in descendant genomes frequently dismantled the syntenic blocks involved in ancient transfers.

As many groups of ODT events that individually appeared as convergent were factorized into unique co-events, the relative frequency of event types that were estimated dramatically changed: relatively to scenarios inferred using a parsimony criterion (minimization of losses) independently applied to single gene histories, block event scenarios resulted in a decrease of 13,421 ODT events, most of them transfer (T) events (10,978, −25.4%), and an increase of loss (L) events (2,896, +9.7%) (Table 2). However, the count of additional losses was certainly over-estimated, because block events of gene loss are bound to have occurred, but we did not intend to factorize loss events in this study (see Methods).

This difference in the estimated number of gene losses was due to the frequent under-estimation of the event age when considering only scenarios for individual gene families, relatively to joint scenarios for several gene families. Indeed, the loss scenarios were generally estimated by fixing the timing of the preceding gene gain (O, D or T) events to their most recent possible location – the most parsimonious solution with respect to losses. In the case of block event scenarios, ODT events were dated to the most recent *common* location of all single-gene event parts, which by definition must be equally ancient as, or more ancient than the single-gene estimates. This resulted in globally older ancestor for block gain events, with a higher number of lineages between the ancestor and extant representatives in which to invoke subsequent losses (Fig. 1 A). ODT events are thought to be less frequent than gene loss (L) events, and the more complex pattern of transfers (characterized by a donor and a recipient) makes it less likely for T events with similar coordinates to have occurred convergently in neighbour genes in the absence of a linkage hypothesis. As a consequence, factorizing similar ODT events for neighbour genes appears a to be a suitable approach to obtain a pangenome-wide scenario that is much more parsimonious in the total number of all kinds of events, i.e. ODTL events.

### Inferred genome histories suggest selection for new genes in ancestors of key *At* lineages

Our inferred history of gain and loss in ancestral genomes of *At* showed heterogeneous dynamics across the species tree. First, the estimated genome sizes were significantly lower in estimated ancestral genomes than in extant genomes (Fig. 2, Table S5). For instance, the estimated genome gene content, or gene repertoire, of the *At* clade ancestor contained around 4,500 genes, while extant genomes had an average size of 5,500 genes. This 1,000-gene difference approximately corresponds to the number of genes recently gained along the terminal branches of the species tree (Fig. 2), indicating a divide in contemporary genomes between a long-standing gene repertoire and a large fraction of newly acquired genes still segregating in the population. Our ancestral genome estimation procedure did not estimate the count of unobserved ancient genes; however, a similar-size polymorphic gene repertoire probably existed in the *At* clade ancestors and was mostly lost in all sampled descendants.

**Figure 2:**
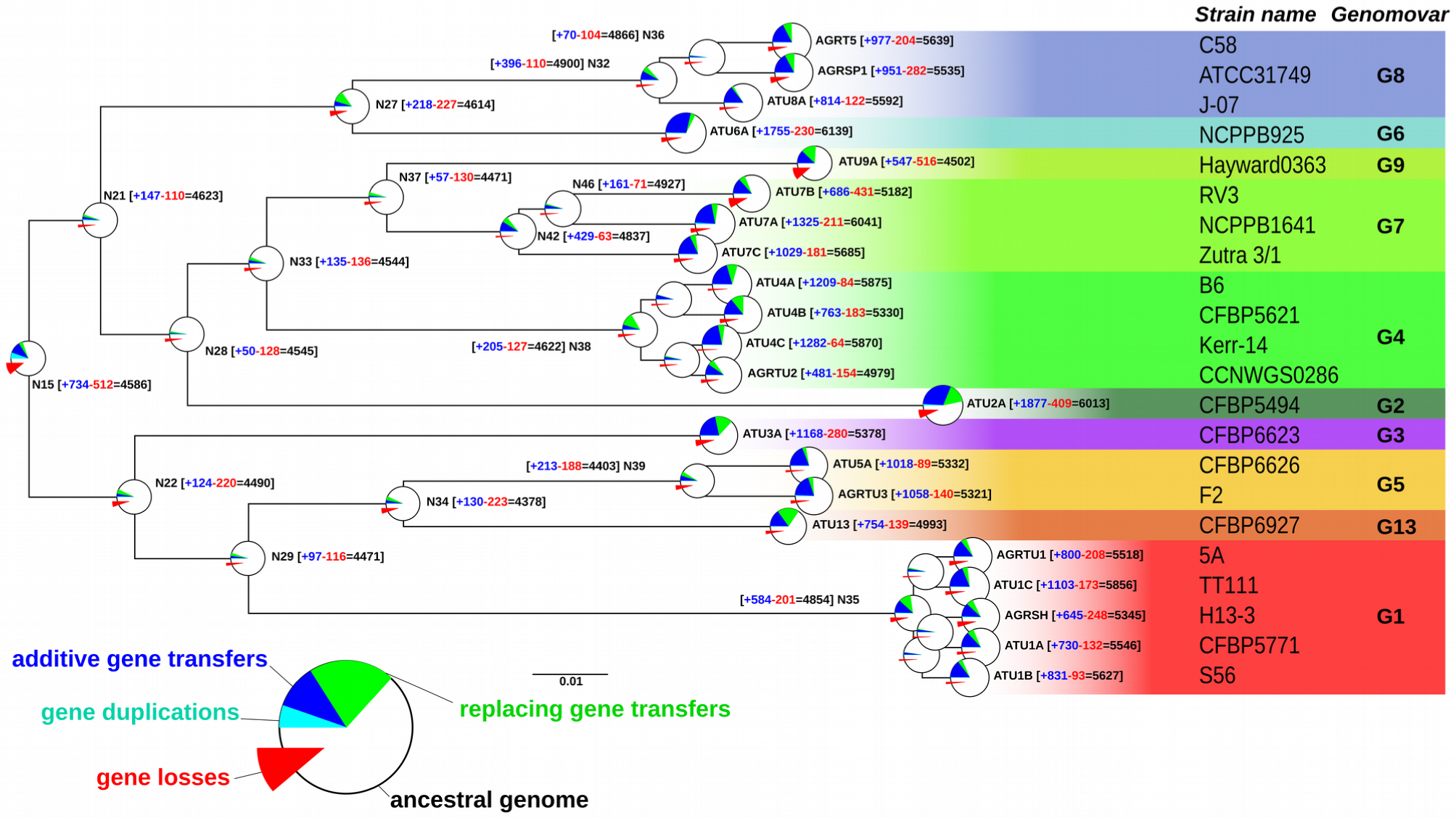
Ancestral genome sizes and gain/loss events. The tree is a subtree of that presented in Fig. S1, and focuses on the *At* clade. Net gains (+) and losses (-) and resulting genome sizes (=) are indicated next to nodes. Disc at inner and terminal nodes represent estimated ancestral genomes and extant genomes, respectively; surfaces are proportional to genome sizes. Prevalence of events shaping the gene content are indicated by pie charts indicating the fraction of losses (red), gains by duplication (cyan), gains by transfer (blue) and gene conversions/allelic replacements (green). The relatively high number of event occurring at the *At* root is related to the long branch from which it stems in the complete Rhizobiales tree (Fig. S1), which is not represented here.

The length of the branch leading to the ancestor best explained the number of genes gained and lost by an ancestor (linear regression, *r*^2^ = 0.59 and 0.32 for gains and losses, respectively), although removing the extreme point of node N35 (the G1 species ancestor) sharply decreased the correlation coefficients (*r*^2^ = 0.27 and 0.28) (Fig. S9 A, B). Interestingly, the number of genes gained by an ancestor and subsequently conserved in all members of the descendant clade, i.e. clade-specific genes, was robustly explained by the ancestor age (*r*^2^ = 0.39, or 0.41 when removing N35) (Fig. S9 F). This relationship was better described by a decreasing exponential regression (*r*^2^ = 0.51, or 0.50 when removing N35), which reflected a process of ‘gene survival’ in genomes over time (Fig. 3). Alternatively, these trends may have resulted from a systematic bias in our estimation procedure: for instance, because our block event inference algorithm tended to place gene gains higher in the species tree than an inference considering a gene family alone would have done (Fig. 1 A), subsequent losses may have been inferred too frequently in early ancestors, generating this pattern of decay over time; however similar trends were observed for scenarios without block aggregation (data not shown).

**Figure 3:**
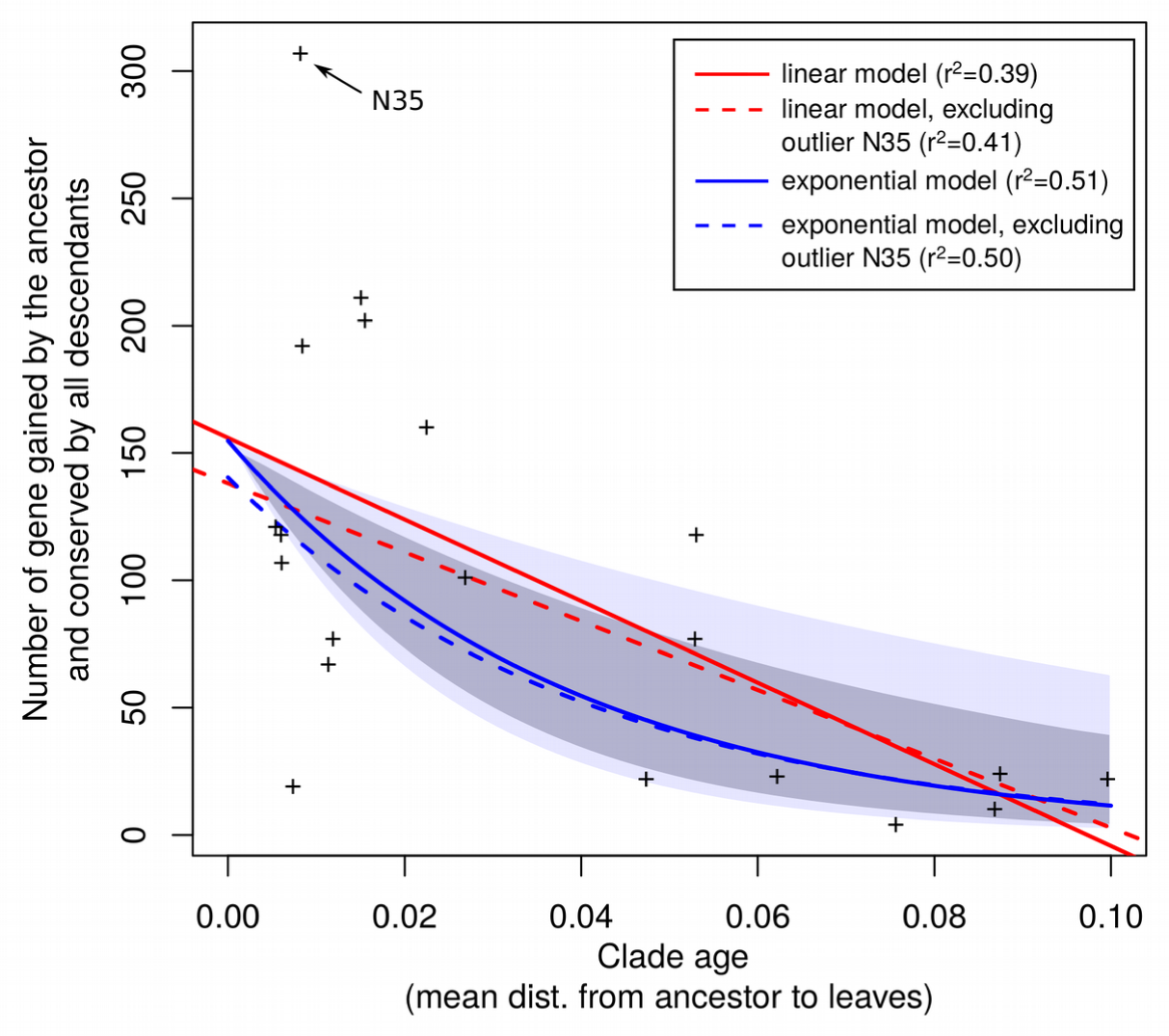
Retention of gained genes within *At* genomes follows a survival model. Node ‘N15’ (the G1 species ancestor) is the strongest driver in the linear regression. Dark and light shaded areas represent the 95% and 99% confidence intervals of the exponential model, respectively (solid blue line).

We identified outlier genomes in this putative ‘gene survival’ process, as the nodes with the largest residuals in the exponential regression (out of the 95% confidence interval). They were, in a decreasing order of excess of conservation relative to their age, the ancestors of the [G6-G8], G1, G5, [G5-G13], G8 clades and those of subclades of G4 and G7 (Fig. S9 F; Fig. S10). These excesses of conservation did not systematically reflect a particular excess of gains in the ancestors: ancestors of G1 and G8 (nodes N35 and N32) did indeed gain more genes than predicted by their respective branch lengths, whereas ancestors of [G6-G8], [G5-G13] and G5 (nodes N27, N34 and N39, respectively) rather lost genes in excess (Fig. S9 C, D). In the latter cases, excess conserved gains may thus have stemmed from a fixation bias like natural selection for new genes. The outliers that fell above this trend – those clades that conserved more genes than predicted by their age – all belonged to [G1-G5-G13] and [G6-G8] clades (Fig. S10). The higher rate of conservation in these clades suggests a higher proportion of genes under purifying selection since their ancestral acquisitions, i.e. domesticated genes.

Clade-specific genes conserved for a long time likely provide a strong adaptive feature to their host organism. A new adaptive trait can improve an organism’s fitness by increasing the differentiation of its ecological niche relatively to cognate species, and thus enable it to escape competition. This emergence of a new ecotype – an ecologically differentiated lineage – can for instance occur through a gain of function (e.g. via additive HGT) that allows for exclusive consumption of a resource (Lassalle et al. 2015) or the change in relative reliance on a set of resources (Kopac et al. 2014). The spread of such niche-specifying traits to close relatives of the ecotype should be counter-selected (Cohan & Koeppel 2008), so that their occurrence is expected to be restricted to the descendants of the ecotype, i.e. to be clade-specific. Identifying such adaptive traits among clade-specific genes is thus the key to the understanding of the unique ecological properties of a bacterial clade.

### Clusters of clade-specific genes are under purifying selection for their collective function

Niche-specifying traits are expected to provide higher differential fitness if they are less likely to be already present in, or independently acquired by, competing relatives. Hence, the best candidates for niche-specifying traits consist of novel and complex traits relying on an array of biochemical functions coded by separate evolving units (genes) and do not depend on pre-existing pathways, making such mutations unlikely to occur several times by chance. In such a case, it is crucial for the complete set of underlying biochemical functions to be gained at once for it to provide any kind of advantage. Such an event can typically happen with the co-transfer of a complete operon. In a previous study focused on G8 genomes (Lassalle et al. 2011), we observed that clade-specific genes tended to occur in clusters of genes with related biochemical function. This apparently non-random pattern of gene conservation suggests that co-transferred groups of genes collectively coding for a function were selected among incoming transferred genes: initially by positive selection for their new function upon transfer reception, and later on by negative (purifying) selection against the destruction of the group by rearrangement or partial deletion. This led us to consider clusters of co-functioning clade-specific genes as good candidates for niche-specifying determinants (Lassalle et al. 2011). Yet, it is well known that bacterial genomes are organized in functional units such as operons, super-operons, etc. (Rocha 2008), and the co-transfer of cooperating genes could neutrally result from the functional structure of the donor genomes. However, the transferred DNA segments are most probably taken randomly from donor genomes, apart from the special case of genes encoding their own mobility. Thus, under a neutral model, co-transferred genes should not always be co-functioning, and the probability for a transferred fragment to span a functional element like an operon is expected to be close to that of any similarly sized fragment of the donor genome.

To test whether clustering of functionally related clade-specific genes resulted from natural selection, we designed tests that assessed the relationship between gene transfer history and functional homogeneity (*FH*) (see Methods). First, we verified that random groups made of physically distant genes had lower *FH* values than groups of neighbour genes, confirming that *FH* captures the functional structure of a genome (Fig. 4 A). Then we compared random groups of neighbour genes without a shared transfer history to blocks of co-transferred genes of the same size. The distribution of *FH* values showed that while blocks of co-transferred genes generally gathered genes that do not encode related functions or for which functional annotations are insufficient (FH ~ 0), a minor fraction presented intermediate to high functional relatedness (e.g. in the G4-B6 genome, minor modes at *FH* ~ 0.35 and *FH* ~ 0.75, Fig. 4 A). Blocks of co-transferred genes had significantly higher *FH* values than random groups in 45 out of 49 significant tests performed on independent combinations of genomes and block sizes (Fig. 4 A, C). This shows that fixation of transferred blocks of genes in genomes was biased towards blocks that code for functional partners in a biological process. This observation supports the hypothesis of positive selection favouring fixation in a recipient genome of the transferred genes that can immediately provide a selectable function. It is also compatible with the ‘selfish operon’ model proposed by Lawrence and Roth (1996): in host genomes, transfer followed by selection for readily functional multi-genic traits is thought to lead to the prevalence of genes clustered into tightly linked functional units.

**Figure 4:**
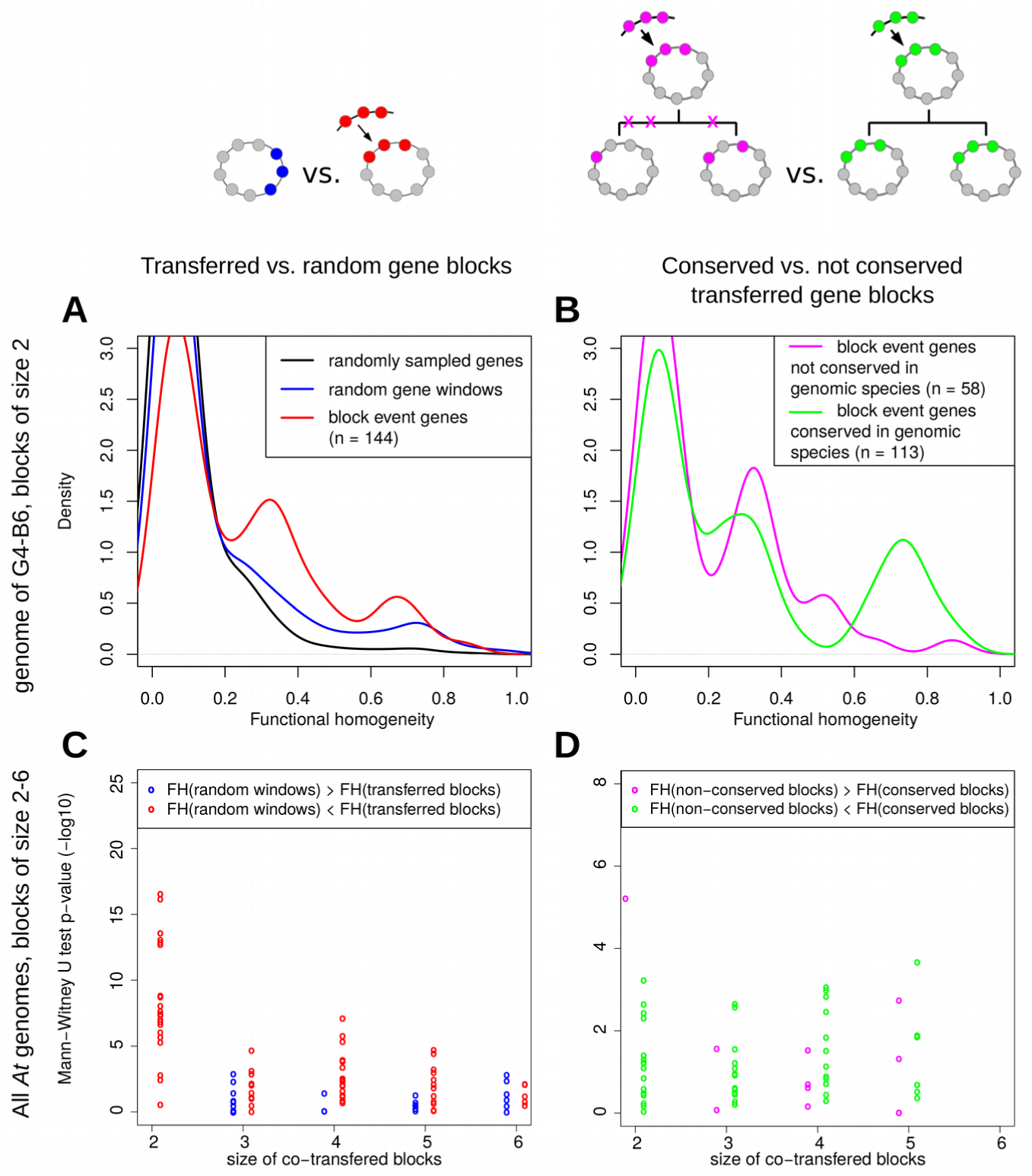
Functional homogeneity of gene clusters. (A, B) Distribution of functional homogeneity (*FH*) values of genes within clusters using representative plots comparing clusters of two genes in the B6 genome (a G4 member). (A) Comparison of *FH* values of groups of two genes taken from the B6 genome: randomly distant pairs (black), any pair of neighbour genes without a common transfer history (blue), or pairs of co-transferred neighbour genes (red). (B) Comparison of *FH* values of pairs of co-transferred genes from families conserved across all G4 strains (green) or not conserved (red). (C, D) Distribution of *p*-values of Mann-Whitney-Wilcoxon sum of ranks test comparing the distributions of *FH values* (made independently for all *At* genomes at all discrete block sizes) of (C) random windows of non co-transferred genes vs. blocks of co-transferred genes or (D) conserved vs. non-conserved blocks of co-transferred genes. Each point represents an observation from an extant *At* genome for a given gene group size (on the *x-*axis). Point colours indicate the higher-*FH* category: (C) blue, *FH*(random windows) > *FH*(transferred blocks), 29/95 tests (4/49 significant tests); red, *FH*(random windows) < *FH*(transferred blocks), 66/95 (45/49); (D) purple, *FH*(non-conserved blocks) > *FH*(conserved blocks), 11/60 (2/13); green, *FH*(non-conserved blocks) < *FH*(conserved blocks), 49/60 (11/13). Tests were considered significant at *p* < 0.01.

In addition, among the groups of genes acquired by transfer, those that were conserved in all descendants of the recipient ancestors had more coherent annotated functions than the non-conserved ones (11/13 significant tests are positive, Fig. 4 B, D). The hypothesis of conserved co-transferred genes encoding more related functions than non-conserved ones was previously proposed based on manual inspection of the functional relatedness of a few transferred operons in *E. coli* (Homma et al. 2007) or the metabolic flux coupling of spatially clustered transferred genes (from possibly mixed origins) in Gammaproteobacteria (Dilthey & Lercher 2015). The present study presents a first quantitative estimation of functional relatedness within blocks of co-transferred genes, and provides a statistical argument for purifying selection enforcing their collective conservation in genomes. This supports our initial hypothesis that clusters of clade-specific genes participating to a same pathway were more likely to carry sufficient information to encode a new adaptive trait, and had been under continued selection since their acquisition. It follows that the adaptations that characterize the ecological niche of a clade should be revealed by identifying of the genes specifically conserved inside a clade, and notably those grouped in clusters with related functions.

### Identification of clade-specific genes in *A. tumefaciens* key clades

We investigated the histories of gene gain and loss in the clades of *At* to identify the synapomorphic presence/absence of genes in these clades. We used an automated method that recognizes profiles of contrasted gene occurrence among sister clades by spotting ancestral gene gains or losses that resulted in their conserved presence or absence in the descendant clade (see Methods). Doing so, we accounted for convergent gains/losses of orthologous genes in distant clades, notably in cases of a transfer from one clade ancestor to another; this allowed us to evidence the specific sharing of genes between non-sister species of *At*. Listings of clade-specific genes of those key *At* clades can be found in Dataset S1, or can be browsed on the Agrogenom database website http://phylariane.univ-lyon1.fr/db/agrogenom/3/ (Fig. 5). Generally speaking, clade-specific genes were often located in relatively large clusters encoding coherent biochemical functions or pathways, which are summarized in Table S6 and hereafter numbered with the AtSp prefix. Those clade-specific gene clusters often matched transfer or origination block events as estimated above (Dataset S1), although often with limited coverage or with several transfer blocks mapping to a single clade-specific cluster. This suggests that block gain events are likely to cluster at the same loci. Alternatively, it suggests a limitation of our search procedure in the face of the complexity of gene histories, with different patterns of multiple consecutive transfers in different gene families preventing recognition of their common history. Extended description of the noteworthy biochemical functions encoded in these clade-specific gene repertoires can be found in the Supplementary Text (section 6). Species G1, G8, G4 and G7, were represented by several closely related extant genomes, and therefore were particularly amenable for the accurate definition of clade-specific gene repertoires. For these species, chromosomal maps (Fig. S11, S12, S13, S14) show that species-specific genes were unevenly located on the various replicons of *At* genomes, with a bias towards accumulation on the linear chromosome (Lc), and an unexpected presence on the At plasmid (pAt) (Tables S6, S7).

**Figure 5:**
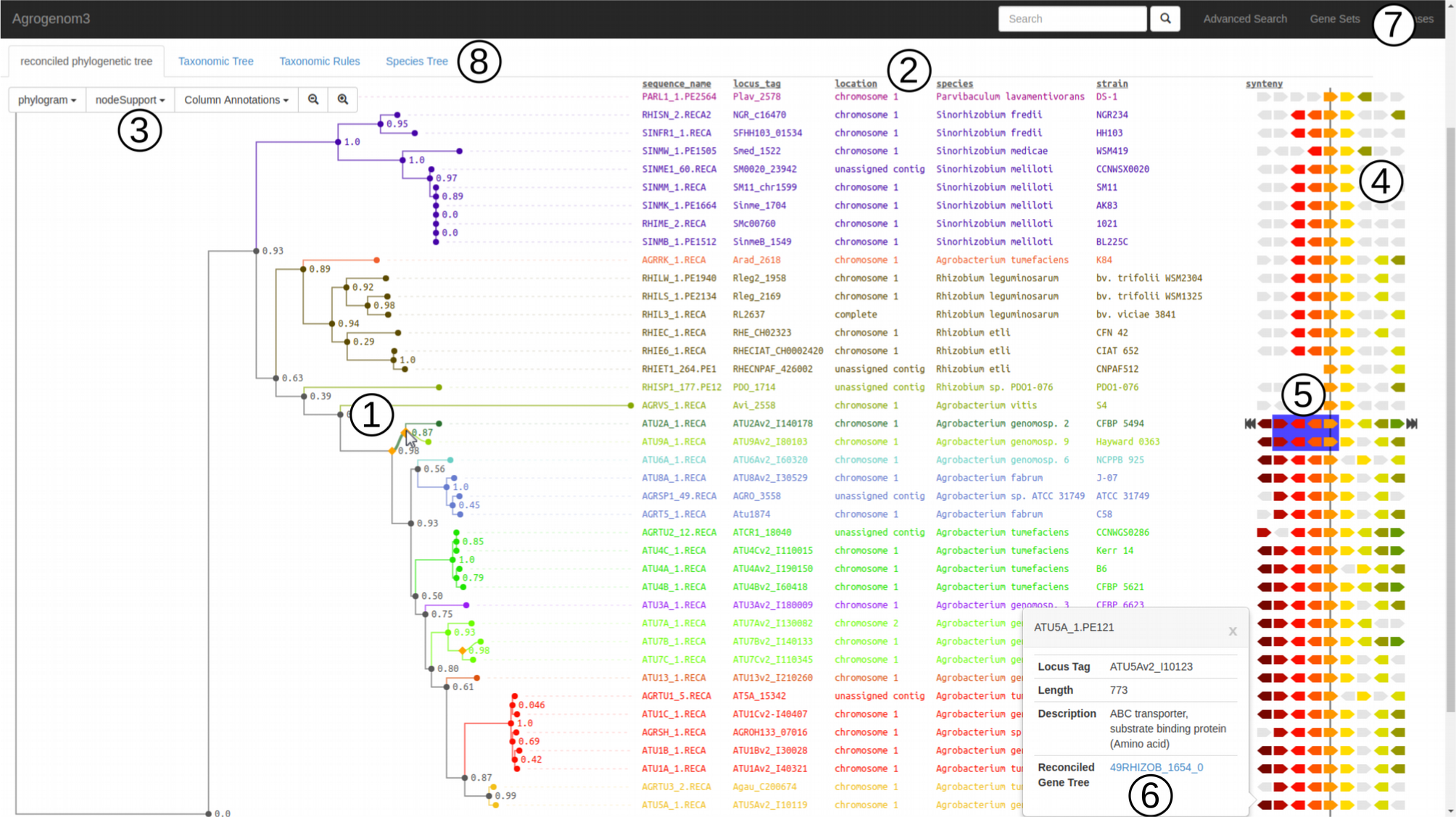
Snapshot of the Agrogenom web interface. View of the *recA* gene family. (1) Reconciled gene tree; the orange diamond under the mouse cursor indicates a transfer event from G2-CFBP 5494 to G9-Hayward 0363. (2) Detailed annotation of the sequences at the tip of the tree, including locus tag (linking out to MaGe genome browser), chromosomal location, taxon name, database cross-references, etc. (3) Dynamic menu to adapt the level of displayed information. (4) Syntenic view in the genomic neighbourhoods of the focal gene family; homologues share the same colour, defined with reference to a chosen sequence (indicated by the navigation arrows on the sides). (5) The blue frame indicates a block transfer event involving four gene families; this block appears dynamically when hovering the cursor above the transfer node in the gene tree. (6) A pop-up window with the functional annotation and characteristics of a gene can be generated by double-clicking on the gene; it contains the link to the gene tree of the gene family. (7) Search menus: rapid search using gene names; ‘Advanced search’ to reach a gene family from its various annotation fields; ‘Gene Sets’ to browse lists of genes: clade-specific genes, core genome, ancestral gene content, clade-specific gains/losses. (8) Alternative views: the reference species tree and a projection of the gene family distribution among taxa.

### Secondary replicons of Agrobacterium genomes bear clade-specific innovations

Rhizobiaceae have complex genomic architectures composed of a primary chromosome, plus a secondary chromosome or megaplasmid bearing essential genes, called the chromid (Harrison et al. 2010), and a variable complement of plasmids of various sizes (Young et al. 2006). More specifically, the chromid of the *Agrobacterium* genus (Mousavi et al. 2015; Ormeño-Orrillo et al. 2015), which includes the *At* clade, is linear (Slater et al. 2009, 2013) as a result of a unique ancestral event of linearization and thus constitutes a synapomorphy of this clade (Ramírez-Bahena et al. 2014). Another general feature of *At* genomes is the frequent presence of a pAt, a megaplasmid that was long referred to as the cryptic plasmid because its role in agrobacterial cell biology remains largely unknown. We found that different pAt types were restricted to certain genomic backgrounds (based on their replication gene phylogenies) and carried clade-specific gene clusters at the species level (in G1, G8, G4 and G7 species) or higher (in [G6-G8] clade) (Fig. S11, S12, S13, S14; Supplementary Text, section 8). pAts therefore appear as core replicons of a majority of *At* species. In addition, while many megaplasmids of the same *repABC* family are known to recombine intensely within species (Kumar et al. 2015; Epstein et al. 2012), the occurrence of clade-specific genes on pAts and never on the other plasmids (pTis and smaller ones) suggests the existence of barriers to its transfer. Within Cohan’s ecotype framework, we interpret this pattern as the presence of determinants of the species’ ecological niche on these particular extra-chromosomal elements, which selectively prevented their spread among closely related species (Cohan & Koeppel 2008). This suggests that the pAt is probably an essential replicon for most species of *At* in their natural environments and qualifies it as a *bona fide* chromid (Harrison et al. 2010). Deletion-mutant competition experiments on the distantly related chromid pSymB (diCenzo et al. 2016) demonstrated that the chromid had a significant regulatory impact on the bacterial host and contribution to its fitness in the plant rhizopshere (i.e. outside of a symbiotic nodule). Consequently, these megaplasmids possibly play an determining role in adaptation to their core ecological niche (Lassalle et al. 2015). Functional investigation of the core functions borne by agrobacterial pAts could thus provide a better understanding of the specific ecophysiology of each *At* species.

### Clade-specific gene functions provide insights into the possible ecological speciation of clade ancestors

The nature of putative ecological specialization is not obvious for agrobacteria, which are ubiquitous soil-dwellers. Different *Agrobacterium* species frequently co-occur in soils, sometimes in the same micro-metric sample (Vogel et al. 2003); based on the competitive exclusion principle (Gause 1932), they must have distinct ecologies. Certain soils and/or host plants are preferentially colonized by certain species (Costechareyre et al. 2010). In parallel, G2 members appear to have developed a capacity towards opportunistic pathogenicity in humans (Aujoulat et al. 2011). This shows some kind of niche differentiation occurs among *Agrobacterium* species, but the precise nature of the underlying environmental factors still remains to be deciphered. Because clade-specific genes are expected to encode what makes the ecology of a clade to be distinct from that of its relatives (Lassalle et al. 2015), we investigated the specific functional repertoire of At clades. Strikingly, in most clades, including species or higher-level groups, the sets of clade-specific genes recurrently presented the same classes of functions. These include transport and metabolism of phenolic compounds, aminoacids and complex sugars, and production of exopolysaccharides and siderophores, all of which can be related to bacterial life in the plant rhizosphere (Lassalle et al. 2011).

Among these, we can notably report the specific presence of a supernumerary chemotaxis regulation operon *che2* in species G1, which is uniquely linked to an array of genes with predicted functions involved in the catabolism of (possibly aminated) aromatic compounds (Table S6). This suggests that G1 strains are able to specifically degrade certain – yet unknown – aromatic compounds, for which they might display specific tropism and/or induction of biofilm formation.

G8 species and the [G6-G8] clade presented a number of clade-specific gene clusters (Table S6), as previously reported (Lassalle et al. 2011), among which the largest were the ferulic acid degradation and siderophore biosynthesis operons. These operons have been reported to provide a growth advantage and to be expressed in a coordinated manner in a plant rhizosphere environment (Campillo et al. 2014; Baude et al. 2016). Taken together, these results show that G8 lineage-specific genes jointly participate in the adaptation to a plant-related specific ecological niche. Interestingly, the gain of a siderophore biosynthesis locus in the [G6-G8] clade ancestor coincided with the loss of the locus encoding biosynthesis of another siderophore, agrobactin, otherwise ubiquitous in, and unique to, the *At* clade. This conserved switch to a different pathway for iron scavenging – a crucial function in iron-depleted plant rhizospheres – may provide a competitive advantage with respect to co-occurring agrobacteria.

The [G5-G13] species group specifically presented a phenylacetate degradation pathway operon (Table S6), which biochemical function was demonstrated *in vitro* (Fig. S15). This discovery readily provides us with a specific biochemical identification test for these species, and again hints to the particular affinity of agrobacteria for aromatic compounds likely to be found in plant rhizospheres.

Finally, the large cluster that encodes the nitrate respiration (denitrification) pathway, including the *nir*, *nor*, *nnr* and *nap* operons was absent from the [G1-G5-G13] clade. More recently, that gene cluster was also lost by strains G9-NCPPB925 and G8-ATCC31749, and its presence in strain G3-CFBP6623 seems to result from later transfer from a mosaic of sources within *At*. Considering the absence of this super-operon in close relatives of *At* such as *A. vitis* and *R. leguminosarum*, it was likely acquired by the ancestor of the [G2-G4-G7-G9-G6-G8] clade (node N21 on Fig. 1), one of the two large clades that divide the *At* complex. Strains possessing the denitrification pathway may be selectively advantaged under certain anaerobic or micro-aerophilic conditions, like those met in certain soils and rhizospheres; such an adaptation may have supported an early differentiation of *At* lineages towards the colonization of partitioned niches.

Species G1 and G8 presented a particular case of convergence of their clade-specific functional repertoire. Firstly, they shared 57 synapomorphic genes (Table S7 and S8), in most cases with phylogenetic support for transfer events among respective ancestors. These traits were previously hypothesized to provide key adaptation to life in the plant rhizosphere of G8 (= *A. fabrum)* (Lassalle et al. 2011). For instance, these species share homologous genes involved in the biosynthesis of curdlan – a cellulose-like polysaccharide – and the biosynthesis of O-antigens of the lipopolysaccharide (LPS) (Table S6; Supplementary Text, section 5.1). These two capsular components may define attachment properties of the cell to the external environment, possibly in a similar way than the LPS synthesized by homologous enzymes in *Brucella spp*., which mediates a specific interaction with cells of a eukaryotic host (Vizcaíno et al. 2001). In addition, nonhomologous G1 and G8 clade-specific genes encoded similar functional pathways, i.e. phenolic compound metabolism and exopolysaccharide production (Table S6).

This convergence of the niche-specifying gene repertoires of species G1 and G8 may have caused a stronger overlap of their ecological niches, which in turn might have led to inter-species competition for resources. However, shared niche-specifying genes occur in combination to different sets of species-specific genes in the core-genome of each species, and different epistatic interactions could induce strong divergence in their phenotype. Typically, even though the loci for LPS O-antigen biosynthesis in G1 and G8 are highly similar (>93% amino acid identity in average for proteins of the homologous AtSp14 loci, Fig. S16) and most likely produce a structurally equivalent compound, regulation of biofilm production by these species is probably different. Indeed, several regulatory genes specific to the G1 genomes are involved in the regulation of chemotaxis/biofilm production, such as the *che2* operon (cluster AtSp2) and hub signal-transducing protein HHSS (“hybrid-hybrid” signal-sensing, see Supplementary Text, section 5.1) found in cluster AtSp14 (Fig. S11 and S16), and a sensor protein (cluster AtSp3) modulating c-di-GMP – a secondary messenger involved in the switch from motile to sessile behaviours. Those specific regulators were all in close linkage to G1-specific genes involved in phenolics catabolism or biofilm production. These latter genes may be the downstream regulatory targets of what seems to be a coherent regulation network controlling motility, biofilm production and phenolics degradation; this locus is potentially coding for a whole pathway for responses to specific environmental conditions of the niche of G1, such as the availability of phenolics to use as nutrients. Similarly, G8-specific genes of the AtSp26 cluster (Fig. S12) formed a regulatory island involved in the perception and transduction of environmental signals, including mechanosensitive channels and a receptor for phenolic compound related to toluene (Lassalle et al. 2011).

Both the G1 and G8 species are thus likely to orchestrate the production of similar polysaccharides under different regulation schemes, involving the coordination of their expression with other specific traits – in both cases the catabolism of (likely different) phenolics. Similarly, coordinated expression of several clade-specific genes resulting in conditional phenotypes has recently been observed in G8-C58 (Baude et al. 2016), strengthening the idea of the existence of an ecological niche to which species G8 is specifically adapted through the expression of a particular *combination* of clade-specific genes. The partial hybridization of the G1- and G8-specific genomes probably led each species to tap the same resources in different ways, avoiding any significant competition between them. These species may thus form guilds of relatives that exploit partitions of a largely common ecological niche (Lassalle et al. 2015), enabling them to co-occur in soils (Vogel et al. 2003; Portier et al. 2006).

While such evolutionary mechanisms of late hybridization and re-assortment of niche-specifying genes have previously been observed (Sheppard et al. 2013), it is unclear whether they are common among other soil/rhizosphere-dwelling bacteria. A recent investigation of the pangenome diversity of *R. leguminosarum* genomic species revealed similar patterns of occurrence of species-specific genes, but none could be related to a species-specific metabolic or symbiotic property, challenging the notion that species could have specific ecological adaptations (Kumar et al. 2015). However, this study only relied on the analysis of the pattern of homologous gene presence/absence, not their gain history, and could have overlooked parallel synapomorphic gene gains. Using our estimation of scenarios of gene evolution, we see that convergent evolution was important in shaping *At* genomes (Table S7, S8) and that ecological niche differentiation may occur through finer processes, including specific regulation of complex sets of functions.

## Conclusion

We developed an original method to estimate the history of all genes in a bacterial pangenome and applied it to the *Agrobacterium* biovar 1 species complex (*At*) to unveil the gain and loss dynamics of the gene repertoire in this taxon. Genes specifically gained by major *At* clades were mostly organized in large blocks of co-evolving genes that encode coherent pathways. This pattern constitutes a departure from a neutral model of gene transfer in bacterial genomes and indicate purifying selection has enforced their conservation. We therefore considered these blocks of clade-specific genes as likely determinants of clade core ecologies. Genes specific to each species and to the *At* species complex as a whole recurrently encoded functions linked to production of secreted secondary metabolites or extracellular matrix, and to the metabolism of plant-derived compounds such as phenolics, sugars and amino acids. These clade-specific genes probably represent parallel adaptations to life in interaction with host plant roots. This suggests that ecological differentiation of *Agrobacterium* clades occurred through the partitioning of ecological resources available in plant rhizospheres. In the future, sampling of within-species diversity, coupled with population genomics approaches, could further reveal ecological properties of agrobacteria, including those that may be non-ubiquitous but dynamically maintained by recombination within species (Kashtan et al. 2014; Rosen et al. 2015). Gene co-evolution models, such as the one developed here, could be extended to the investigation of inter-locus linkage in genome populations (Cui et al. 2015). Such analyses could reveal complex interactions between molecular pathways under ecological selection, opening onto new steps towards the understanding of bacterial adaptation to the infinite diversity of micro-environments.

## Acknowledgements

This project was supported by the French National Research Agency (ANR, http://www.agence-nationalerecherche.fr/) grant ECOGENOME (ANR-BLAN-08-0090) and ANCESTROME (ANR-10-BINF-01-01) and by AGROMICS grant from the ENVIROMICS challenge of the Interdisciplinary Mission of the French National Centre for Scientific Research (CNRS). This work was performed using the computing facilities of the CC LBBE/PRABI. We thank the LABGeM (CEA/IG/Genoscope & CNRS UMR8030) and the France Génomique National infrastructure (funded as part of the ‘Programme Investissement d’Avenir’ of the ANR, contract ANR-10-INBS-09) for support within the MicroScope annotation platform. We thank the associate editor David Bryant and two anonymous reviewers for their thoughtful comments on the manuscript, and Annie Buchwalter for her careful correction of English language.

## Availability of supporting data

The sixteen new genome sequences used in these projects were submitted to the EBI-ENA (www.ebi.ac.uk/ena) under the BioProjects PRJEB12180-PRJEB12196. Accession numbers for the corresponding replicon sequences are LN999991-LN999996, LT009714-LT009764 and LT009775-LT009780. Accession numbers of all genomic data used in this dataset are listed Table 1. All other relevant data (output of analyses) are available on Figshare at https://figshare.com/projects/Ancestral_genome_reconstruction_reveals_the_history_of_ecological_diversification_in_Agrobacterium/20894. The original software tools developed for this work are available at https://github.com/flass/agrogenom.

## Competing interests

The authors declare no competing interests about the present study.

## Authors’ contributions

FL, XN and VD conceived and supervised the study. DC, DMu and LV cultivated the bacteria, prepared the samples and conducted biochemical tests. FL, RP, SP, TB and LG contributed to coding and developing software. FL and RP designed the Agrogenom database. RP designed the Agrogenom web interface. DMo generated functional annotation data. AC and DV integrated data into the Microscope database. VB led the sequencing project. VB, FL and AD participated in genome assembly. FL, TB and VD conceived the phylogenetic methods. FL implemented the phylogenetic methods and performed the statistical and genomic analyses. FL, LV, DMu, VD and XN participated in writing the manuscript.

## List of abbreviations

*At*: *Agrobacterium tumefaciens* species complex

Cc: circular chromosomes

CDS: coding sequence

*FH*: functional homogeneity

HGT: horizontal gene transfer

HHSS: “hybrid-hybrid” signal-sensing protein

Lc: linear chromid

ODT: origination, duplication and transfer (events)

ODTL: origination, duplication, transfer and loss (events)

pAt: *At* plasmid

pTi: Tumor-inducing plasmid

## Supporting Information

**Figure S1. Phylogeny of 131 Alpha-proteobacteria genomes.**

**Figure S2. Reference phylogeny of Rhizobiales history.**

**Figure S3. Bioinformatic pipeline for reconciliation of gene and genome histories.**

**Figure S4. Algorithm for construction of blocks of co-transferred genes.**

**Figure S5. Alternative reference tree topologies obtained with different methods.**

**Figure S6. Support for monophyly of groups in all gene trees.**

**Figure S7. Hierarchical clustering of *Rhizobiales* genomes according to their gene content.**

**Figure S8. Distribution of block event sizes.**

**Figure S9. Gene gain, loss and conservation within *At* clade ancestors.**

**Figure S10. Residuals of the negative exponential regression of clade age vs. conservation of gained genes.**

**Figure S11. Historical stratification of gains in the lineage of *A. sp* G1 strain TT111.**

**Figure S12. Historical stratification of gains in the lineage of *A. sp.* G8 (*A. fabrum*) strain C58.**

**Figure S13. Historical stratification of gains in the lineage of *A. sp* G4 (*A. radiobacter*) strain B6.**

**Figure S14. Historical stratification of gains in the lineage of *A. sp. G7* strain Zutra 3/1.**

**Figure S15. Growth curves of representative of *At* genomic species on phenylacetate.**

**Figure S16. Syntenic conservation of the AtSp14 cluster in G1, G8 and Brucelaceae.**

**Table S1. Statistics of the 16 new genome sequences.**

**Table S2. List of the 455 universal unicopy gene families.**

**Table S3. Matrix of the presence/absence of the 49 ribosomal gene families in the 47 Rhizobiaceae genomes.**

**Table S4. Bioinformatic pipeline for homologous database construction and gene tree/species tree reconciliation.**

**Table S5. Statistics of gains and losses per contemporary and ancestral genome and replicon.**

**Table S6. Location and functional description of clade-specific gene clusters in *A. tumefaciens* genomes.**

**Table S7. Summary of clade-specific genes in the TT111 genome.**

**Table S8. Summary of clade-specific genes in the C58 genome.**

**Supplementary Text. Section 1.** Comparison of several hypotheses for the core-genome reference phylogeny. **Section 2.** Construction of the Agrogenom database. **Section 3.** Reconciliation of gene trees with the species tree. **Section 4.** Block event inference: algorithms. **Section 5.** Clade-specific genes: insights into the ecological properties of clades. **Section 6.** Selected cases of large transfer events. **Section 7.** Secondary replicons of *Agrobacterium* genomes bear clade-specific innovations.

**Dataset S1. Lists of clade-specific genes per clade.**

## References

Abby SS, Tannier E, Gouy M, Daubin V. 2010. Detecting lateral gene transfers by statistical reconciliation of phylogenetic forests. BMC Bioinformatics.

Abby SS, Tannier E, Gouy M, Daubin V. 2012. Lateral gene transfer as a support for the tree of life. Proc. Natl. Acad. Sci. 109:4962–4967.

Ashburner M et al. 2000. Gene Ontology: tool for the unification of biology. Nat. Genet. 25:25–29.

Aujoulat F et al. 2011. Multilocus sequence-based analysis delineates a clonal population of *Agrobacterium* (*Rhizobium*) *radiobacter* (*Agrobacterium tumefaciens*) of human origin. J. Bacteriol. 193:2608–2618.

Baude J et al. 2016. Coordinated Regulation of Species-Specific Hydroxycinnamic Acid Degradation and Siderophore Biosynthesis Pathways in *Agrobacterium fabrum*. Appl. Environ. Microbiol. 82:3515–3524.

Bigot T, Daubin V, Lassalle F, Perrière G. 2013. TPMS: a set of utilities for querying collections of gene trees. BMC Bioinformatics. 14:109.

Bruen TC, Philippe H, Bryant D. 2006. A Simple and Robust Statistical Test for Detecting the Presence of Recombination. Genetics. 172:2665–2681.

Campillo T et al. 2014. Analysis of Hydroxycinnamic Acid Degradation in *Agrobacterium fabrum* Reveals a Coenzyme A-Dependent, Beta-Oxidative Deacetylation Pathway. Appl. Environ. Microbiol. 80:3341–3349.

Cohan FM, Koeppel AF. 2008. The origins of ecological diversity in prokaryotes. Curr. Biol. CB. 18:R1024-1034.

Cooper VS, Vohr SH, Wrocklage SC, Hatcher PJ. 2010. Why Genes Evolve Faster on Secondary Chromosomes in Bacteria. PLoS Comput Biol. 6:e1000732.

Costechareyre D et al. 2010. Rapid and Efficient Identification of *Agrobacterium* Species by recA Allele Analysis : *Agrobacterium* recA Diversity. Microb. Ecol. 60:862–72.

Csűrös M. 2008. Ancestral Reconstruction by Asymmetric Wagner Parsimony over Continuous Characters and Squared Parsimony over Distributions. In: Comparative Genomics. Nelson, CE & Vialette, S, editors. Lecture Notes in Computer Science Springer Berlin Heidelberg pp. 72–86.

Cui Y et al. 2015. Epidemic Clones, Oceanic Gene Pools, and Eco-LD in the Free Living Marine Pathogen *Vibrio parahaemolyticus*. Mol. Biol. Evol. 32:1396–1410.

Daubin V, Lerat E, Perrière G. 2003. The source of laterally transferred genes in bacterial genomes. Genome Biol. 4:R57.

David LA, Alm EJ. 2011. Rapid evolutionary innovation during an Archaean genetic expansion. Nature. 469:93–96.

Dimmer EC et al. 2011. The UniProt-GO Annotation database in 2011. Nucleic Acids Res. 40:D565–D570.

Doolittle WF. 2013. Is junk DNA bunk? A critique of ENCODE. Proc. Natl. Acad. Sci. 110:5294–5300.

Doyon J-P, Ranwez V, Daubin V, Berry V. 2011. Models, Algorithms and Programs for Phylogeny Reconciliation. Brief. Bioinform. 12:392–400.

Felsenstein J. 1993. PHYLIP (Phylogeny Inference Package) version 3.5c.

Fitch WM. 1971. Toward Defining the Course of Evolution: Minimum Change for a Specific Tree Topology. Syst. Zool. 20:406.

Friedman J, Alm EJ, Shapiro BJ. 2013. Sympatric Speciation: When Is It Possible in Bacteria? PLoS ONE. 8:e53539.

Gause GF. 1932. Experimental Studies on the Struggle for Existence I. Mixed Population of Two Species of Yeast. J. Exp. Biol. 9:389–402.

Gonzalez V et al. 2010. Conserved Symbiotic Plasmid DNA Sequences in the Multireplicon Pangenomic Structure of *Rhizobium etli*. Appl Env. Microbiol. 76:1604–1614.

González V et al. 2003. The mosaic structure of the symbiotic plasmid of *Rhizobium etli* CFN42 and its relation to other symbiotic genome compartments. Genome Biol. 4:R36–R36.

Goodner B et al. 2001. Genome sequence of the plant pathogen and biotechnology agent Agrobacterium tumefaciens C58. Science. 294:2323–2328.

Guindon S, Gascuel O. 2003. A Simple, Fast, and Accurate Algorithm to Estimate Large Phylogenies by Maximum Likelihood. Syst. Biol. 52:696–704.

Hao X, Xie P, et al. 2012. Genome Sequence and Mutational Analysis of Plant-Growth-Promoting Bacterium *Agrobacterium tumefaciens* CCNWGS0286 Isolated from a Zinc-Lead Mine Tailing. Appl. Environ. Microbiol. 78:5384–5394.

Hao X, Lin Y, et al. 2012. Genome Sequence of the Arsenite-Oxidizing Strain *Agrobacterium tumefaciens* 5A. J. Bacteriol. 194:903.

Harrison PW, Lower RPJ, Kim NKD, Young JPW. 2010. Introducing the bacterial ‘chromid’: not a chromosome, not a plasmid. Trends Microbiol. 18:141–148.

Homma K, Fukuchi S, Nakamura Y, Gojobori T, Nishikawa K. 2007. Gene Cluster Analysis Method Identifies Horizontally Transferred Genes with High Reliability and Indicates that They Provide the Main Mechanism of Operon Gain in 8 Species of Gamma-Proteobacteria. Mol. Biol. Evol. 24:805–813.

Kashtan N et al. 2014. Single-Cell Genomics Reveals Hundreds of Coexisting Subpopulations in Wild *Prochlorococcus*. Science. 344:416–420.

Kettler GC et al. 2007. Patterns and Implications of Gene Gain and Loss in the Evolution of Prochlorococcus. PLoS Genet. 3:e231.

Kopac S et al. 2014. Genomic Heterogeneity and Ecological Speciation within One Subspecies of *Bacillus subtilis*. Appl. Environ. Microbiol. 80:4842–4853.

Kumar N et al. 2015. Bacterial genospecies that are not ecologically coherent: population genomics of *Rhizobium leguminosarum*. Open Biol. 5:140133.

Kuo C-H, Moran NA, Ochman H. 2009. The consequences of genetic drift for bacterial genome complexity. Genome Res. 19:1450–1454.

Kurtz S et al. 2004. Versatile and open software for comparing large genomes. Genome Biol. 5:R12.

Lassalle F et al. 2011. Genomic Species Are Ecological Species as Revealed by Comparative Genomics in *Agrobacterium tumefaciens*. Genome Biol. Evol. 3:762–781.

Lassalle F, Muller D, Nesme X. 2015. Ecological speciation in bacteria: reverse ecology approaches reveal the adaptive part of bacterial cladogenesis. Res. Microbiol. 166:729–741.

Lawrence JG, Ochman H. 1997. Amelioration of Bacterial Genomes: Rates of Change and Exchange. J. Mol. Evol. 44:383–397.

Lawrence JG, Roth JR. 1996. Selfish operons: horizontal transfer may drive the evolution of gene clusters. Genetics. 143:1843–1860.

Li A et al. 2011. Genome Sequence of Agrobacterium tumefaciens Strain F2, a Bioflocculant-Producing Bacterium. J. Bacteriol. 193:5531–5531.

Makarova K et al. 2006. Comparative genomics of the lactic acid bacteria. Proc. Natl. Acad. Sci. 103:15611–15616.

Marri PR, Harris LK, Houmiel K, Slater SC, Ochman H. 2008. The Effect of Chromosome Geometry on Genetic Diversity. Genetics. 179:511–516.

Morrow JD, Cooper VS. 2012. Evolutionary Effects of Translocations in Bacterial Genomes. Genome Biol. Evol. 4:1256–1262.

Mousavi SA, Willems A, Nesme X, de Lajudie P, Lindström K. 2015. Revised phylogeny of Rhizobiaceae: Proposal of the delineation of *Pararhizobium* gen. nov., and 13 new species combinations. Syst. Appl. Microbiol. 38:84–90.

Ormeño-Orrillo E et al. 2015. Taxonomy of rhizobia and agrobacteria from the Rhizobiaceae family in light of genomics. Syst. Appl. Microbiol. 38:287–291.

Penel S et al. 2009. Databases of homologous gene families for comparative genomics. BMC Bioinformatics. 10:S3.

Pesquita C, Faria D, Falcão AO, Lord P, Couto FM. 2009. Semantic Similarity in Biomedical Ontologies. PLoS Comput Biol. 5:e1000443.

Portier P et al. 2006. Identification of genomic species in *Agrobacterium* biovar 1 by AFLP genomic markers. Appl. Environ. Microbiol. 72:7123–7131.

Ramírez-Bahena MH et al. 2014. Single acquisition of protelomerase gave rise to speciation of a large and diverse clade within the *Agrobacterium*/*Rhizobium* supercluster characterized by the presence of a linear chromid. Mol. Phylogenet. Evol. 73:202–207.

Rocha EPC. 2008. The Organization of the Bacterial Genome. Annu. Rev. Genet. 42:211–233.

Rosen MJ, Davison M, Bhaya D, Fisher DS. 2015. Fine-scale diversity and extensive recombination in a quasisexual bacterial population occupying a broad niche. Science. 348:1019–1023.

Ruffing AM, Castro-Melchor M, Hu W-S, Chen RR. 2011. Genome sequence of the curdlan-producing *Agrobacterium* sp. strain ATCC 31749. J. Bacteriol. 193:4294–4295.

Schlicker A, Domingues FS, Rahnenführer J, Lengauer T. 2006. A new measure for functional similarity of gene products based on Gene Ontology. BMC Bioinformatics. 7:302.

Sheppard SK et al. 2013. Progressive genome-wide introgression in agricultural Campylobacter coli. Mol. Ecol. 22:1051–1064.

Slater S et al. 2013. Reconciliation of Sequence Data and Updated Annotation of the Genome of Agrobacterium tumefaciens C58, and Distribution of a Linear Chromosome in the Genus *Agrobacterium*. Appl. Environ. Microbiol. 79:1414–1417.

Slater SC et al. 2009. Genome sequences of three *Agrobacterium* biovars help elucidate the evolution of multichromosome genomes in bacteria. J. Bacteriol. 191:2501–2511.

Stamatakis A. 2006. RAxML-VI-HPC: Maximum Likelihood-Based Phylogenetic Analyses with Thousands of Taxa and Mixed Models. Bioinformatics. 22:2688–2690.

Szöllősi GJ, Boussau B, Abby SS, Tannier E, Daubin V. 2012. Phylogenetic modelling of lateral gene transfer reconstructs the pattern and relative timing of speciations. Proc. Natl. Acad. Sci. 109:17513–17518.

Touchon M et al. 2009. Organised genome dynamics in the *Escherichia coli* species results in highly diverse adaptive paths. PLoS Genet. 5:e1000344.

Vallenet D et al. 2009. MicroScope: a platform for microbial genome annotation and comparative genomics. Database J. Biol. Databases Curation. 2009:bap021.

Vallenet D. MicroScope Home - MaGe: Microbial Genome Annotation & Analysis Platform - MicroScope - Web Interface System & Specialized Databases for (re)Annotation and Analysis of Microbial Genomes. MicroScope. https://www.genoscope.cns.fr/agc/microscope/home/index.php (Accessed December 18, 2015).

Vallenet D et al. 2013. MicroScope--an integrated microbial resource for the curation and comparative analysis of genomic and metabolic data. Nucleic Acids Res. 41:D636-647.

Vizcaíno N, Cloeckaert A, Zygmunt MS, Fernández-Lago L. 2001. Characterization of a *Brucella* species 25-kilobase DNA fragment deleted from Brucella abortus reveals a large gene cluster related to the synthesis of a polysaccharide. Infect. Immun. 69:6738–6748.

Vogel J, Normand P, Thioulouse J, Nesme X, Grundmann GL. 2003. Relationship between spatial and genetic distance in Agrobacterium spp. in 1 cubic centimeter of soil. Appl. Environ. Microbiol. 69:1482–1487.

Volff JN, Altenbuchner J. 2000. A new beginning with new ends: linearisation of circular chromosomes during bacterial evolution. FEMS Microbiol. Lett. 186:143–150.

Wibberg D et al. 2011. Complete genome sequencing of *Agrobacterium* sp. H13-3, the former *Rhizobium lupini* H13-3, reveals a tripartite genome consisting of a circular and a linear chromosome and an accessory plasmid but lacking a tumor-inducing Ti-plasmid. J. Biotechnol. In Press, Uncorrected Proof.

Williams D, Gogarten JP, Papke RT. 2012. Quantifying Homologous Replacement of Loci between Haloarchaeal Species. Genome Biol. Evol. 4:1223–1244.

Wood DW et al. 2001. The genome of the natural genetic engineer *Agrobacterium tumefaciens* C58. Science. 294:2317–2323.

Young JPW et al. 2006. The genome of *Rhizobium leguminosarum* has recognizable core and accessory components. Genome Biol. 7:R34.

